# Sex, racial, and *APOE*-ε4 allele differences in longitudinal white matter microstructure in multiple cohorts of aging and Alzheimer’s disease

**DOI:** 10.1101/2024.06.10.598357

**Authors:** Amalia Peterson, Aditi Sathe, Dimitrios Zaras, Yisu Yang, Alaina Durant, Kacie D. Deters, Niranjana Shashikumar, Kimberly R. Pechman, Michael E. Kim, Chenyu Gao, Nazirah Mohd Khairi, Zhiyuan Li, Tianyuan Yao, Yuankai Huo, Logan Dumitrescu, Katherine A. Gifford, Jo Ellen Wilson, Francis Cambronero, Shannon L. Risacher, Lori L. Beason-Held, Yang An, Konstantinos Arfanakis, Guray Erus, Christos Davatzikos, Duygu Tosun, Arthur W. Toga, Paul M. Thompson, Elizabeth C. Mormino, Panpan Zhang, Kurt Schilling, Alzheimer’s Disease Neuroimaging Initiative (ADNI), The BIOCARD Study Team, The Alzheimer’s Disease Sequencing Project (ADSP), Marilyn Albert, Walter Kukull, Sarah A. Biber, Bennett A. Landman, Sterling C. Johnson, Julie Schneider, Lisa L. Barnes, David A. Bennett, Angela L. Jefferson, Susan M. Resnick, Andrew J. Saykin, Timothy J. Hohman, Derek B. Archer

## Abstract

**INTRODUCTION:** The effects of sex, race, and Apolipoprotein E (*APOE*) – Alzheimer’s disease (AD) risk factors – on white matter integrity are not well characterized.

*METHODS:* Diffusion MRI data from nine well-established longitudinal cohorts of aging were free-water (FW)-corrected and harmonized. This dataset included 4,702 participants (age=73.06 ± 9.75) with 9,671 imaging sessions over time. FW and FW-corrected fractional anisotropy (FA_FWcorr_) were used to assess differences in white matter microstructure by sex, race, and *APOE-*ε4 carrier status.

*RESULTS:* Sex differences in FA_FWcorr_ in association and projection tracts, racial differences in FA_FWcorr_ in projection tracts, and *APOE-*ε4 differences in FW limbic and occipital transcallosal tracts were most pronounced.

*DISCUSSION:* There are prominent differences in white matter microstructure by sex, race, and *APOE-* ε4 carrier status. This work adds to our understanding of disparities in AD. Additional work to understand the etiology of these differences is warranted.

*Highlights:*

- Sex, race, and *APOE-*ε4 carrier status relate to white matter microstructural integrity
- Females generally have lower FA_FWcorr_ compared to males
- Non-Hispanic Black adults generally have lower FA_FWcorr_ than non-Hispanic White adults
- *APOE-*ε4 carriers tended to have higher FW than non-carriers

*Research in Context:* 

*Systematic Review:* The authors used PubMed and Google Scholar to review literature that used conventional and free-water (FW)-corrected microstructural metrics to evaluate sex, race, and *APOE-*ε4 differences in white matter microstructure. While studies have previously explored differences by sex and *APOE-*ε4 status, less is known about racial differences and no large-scale FW-corrected analysis has been performed.

*Interpretation:* Sex and race were more associated with FA_FWcorr_ while *APOE-*ε4 status was associated with FW metrics. Association, projection, limbic, and occipital transcallosal tracts showed the greatest differences.

*Future Direction:* Future studies to determine the biological and social pathways that lead to sex, racial, and *APOE-*ε4 differences are warranted.

**Consent Statement:** All participants provided informed consent in their respective cohort studies.

## Background

Although Alzheimer’s disease (AD) is traditionally associated with grey matter pathology, emerging data highlights distinct white matter abnormalities in AD, including axonal loss,^1^ demyelination,^2^ and microglial activation,^3^ that can occur up to 20 years prior to symptom onset.^4^ In familial AD, where amyloid and tau accumulate at a young age and there are few co-occurring pathologies or vascular risk factors,^5^ white matter microstructural damage precedes detectable changes in both hippocampal volume and clinical symptoms.^6^ This suggests that white matter damage in AD is not solely due to microvascular disease, and at least in part relates to the underlying pathological mechanisms driving the development of AD. Females, self-identified non-Hispanic Black adults, and Apolipoprotein E (*APOE*)-ε4 carriers are at greater risk of clinical AD, but the pathways by which this occurs are still being elucidated. Examining how white matter integrity differs in these populations is important for understanding disparities in AD and developing targeted interventions.

Two-thirds of people living with AD are women.^7^ There are striking sex differences in AD risk factors, clinical presentation, and neuropathological burden, though the underlying reasons for these differences are not well understood.^8^ Several studies,^9-11^ but not all,^12-14^ have found that females have worse white matter integrity than males. However, many of these studies are limited by small sample size,^12-14^ cross-sectional design,^12,13^ and use of conventional diffusion magnetic resonance imaging (dMRI),^9,14^ thus limiting our understanding of how the trajectory of white matter integrity in aging differs by sex.

Racial categories are social constructs that serve as proxies for sociocultural forces, including social, economic, and environmental factors, that ultimately affect cognition.^15^ Non-Hispanic Black Americans are twice as likely to have AD and related dementias compared to non-Hispanic White Americans,^7^ but are less likely to have amyloid pathology.^16,17^ This suggests that non-amyloid pathways impacted by social and cultural factors also contribute to cognitive impairment. Previous work shows that white matter integrity is affected by social determinants of health such as socioeconomic status,^18,19^ and that observed racial differences are driven by these factors.^20^ However, there is limited work into which tracts are preferentially vulnerable to degradation and whether sex and *APOE*-ε4 carrier status differentially affect white matter integrity in Black and White adults.

*APOE-*ε4 is the strongest genetic risk factor for late-onset AD.^21^ The association between *APOE*-ε4 and AD risk is stronger in females and non-Hispanic White adults compared to males^22^ and non-Hispanic Black adults.^23^ *APOE* has an important role in cholesterol transport and lipid metabolism and may play a role in myelin maintenance,^24^ but the effect of *APOE*-ε4 on white matter integrity is not well-established.^25,26^

To date, most studies of white matter microstructural integrity use conventional dMRI metrics. Conventional dMRI is derived from a single tensor model and is confounded by partial volume effects, such that each voxel contains both tissue and fluid compartments. Free-water (FW) elimination is an advanced post-processing technique that allows for the separation of fluid (FW) and tissue (FW-corrected) components, correcting for partial volume,^27^ to give a FW-corrected fractional anisotropy (FA_FWcorr_) metric. Importantly, FW metrics are considered more sensitive to abnormal brain aging.^28-30^ While previous studies have found loss of white matter integrity in limbic, association, and transcallosal (TC) tracts^28,29,31-35^ associated with cognitive impairment and AD, there has yet to be a large-scale analysis leveraging FW measures to understand how sex, race, and *APOE-*ε4 carrier status are associated with longitudinal white matter integrity.

The goal of this study is to provide the most comprehensive picture to date of the effects of sex, race, and *APOE-*ε4 carrier status on white matter integrity over the course of aging and AD using nine well-characterized cohorts of older adults. In 4,702 participants, leveraging 9,671 harmonized longitudinal imaging sessions, we hypothesized that females, non-Hispanic Black, and *APOE-*ε4 carrying participants would have worse white matter integrity, and that these differences would be pronounced in limbic, association, and transcallosal tracts.

## Methods

### Participants

The present study used data from participants in the Alzheimer’s Disease Neuroimaging Initiative (ADNI), the Baltimore Longitudinal Study of Aging (BLSA), the Biomarkers of Cognitive Decline Among Normal Individuals Study (BIOCARD), the National Alzheimer’s Coordinating Center (NACC) data set, the Religious Orders Study/Rush Memory and Aging Project/Minority Aging Research Study (ROS/MAP/MARS), Vanderbilt Memory and Aging Project (VMAP), and the Wisconsin Registry of Alzheimer’s Prevention (WRAP) cohorts.

ADNI (www.adni.loni.usc.edu), begun in 2003, was designed to assess brain structure and function using serial MRI, other biological markers, and clinical and neuropsychological assessments.^36^ Three ADNI phases (ADNI-GO, ADNI 2, and ADNI 3) were included in the present study. The BLSA was designed to assess physical and cognitive measures in a community-dwelling cohort.^37^ Behavioral assessments began in 1994 and included dementia-free participants aged 55-85. From 2006 to 2018, BLSA MRI data was collected on a 1.5T scanner and, beginning in 2009, MRIs were performed with a single 3T MRI scanner. Data from the BLSA cohort are available upon request by a proposal submission through the BLSA website (www.blsa.nih.gov). BIOCARD, begun in 1995, includes participants who were middle age and cognitively intact at baseline. The study stopped in 2005 and was reestablished in 2009 with annual assessments.^38^ NACC maintains a database of participant information collected from past and present National Institute on Aging-funded Alzheimer’s Disease Research Centers.^39^ ROS/MAP/MARS are longitudinal, epidemiologic clinical-pathologic cohort studies that were designed to characterize common chronic conditions of aging and the neuropathological basis of cognitive impairment. The Religious Orders Study (ROS) was started in 1994 and enrolled older Catholic priests, nuns, and brothers from states across the United States.^40^ The Rush Memory and Aging Project (MAP) started in 1997 and enrolled older men and women in the Chicagoland area.^40^ The Minority Aging Research Study (MARS) was started in 2004 and enrolled older adults who self-identify as Black in the Chicagoland area.^41^ Notably, the three cohorts are managed by a single team with a large common core of data at the item level with imaging managed through a single pipeline allowing efficient merging of data.^42^ VMAP began in 2012 with the goal of understanding the relationship between vascular and brain health.^43^ WRAP, begun in 2001, is a study of midlife adults enriched with persons with a parental history of AD.^44^ For each cohort, demographic and clinical covariates were required for inclusion, including age, sex, educational attainment, race/ethnicity, *APOE* haplotype status (*ε2, ε3*, ε4), and cognitive diagnosis (cognitively unimpaired, mild cognitive impairment, AD). Each cohort had its own inclusion/exclusion criteria. Participants were included in this study if they had dMRI data, demographic and clinical data, were 50+ years old, identified as non-Hispanic White or Black, and passed neuroimaging quality control procedures. Across all cohorts, written informed consent was provided by participants, and research was conducted in accordance with approved Institutional Review Board (IRB) protocols. Secondary analysis of these data was approved by the Vanderbilt University IRB. **Table 1** provides an overview of the ADNI, BLSA, BIOCARD, NACC, ROS/MAP/MARS, VMAP and WRAP sample sizes, demographic information, and health characteristics.

**Table 1.**
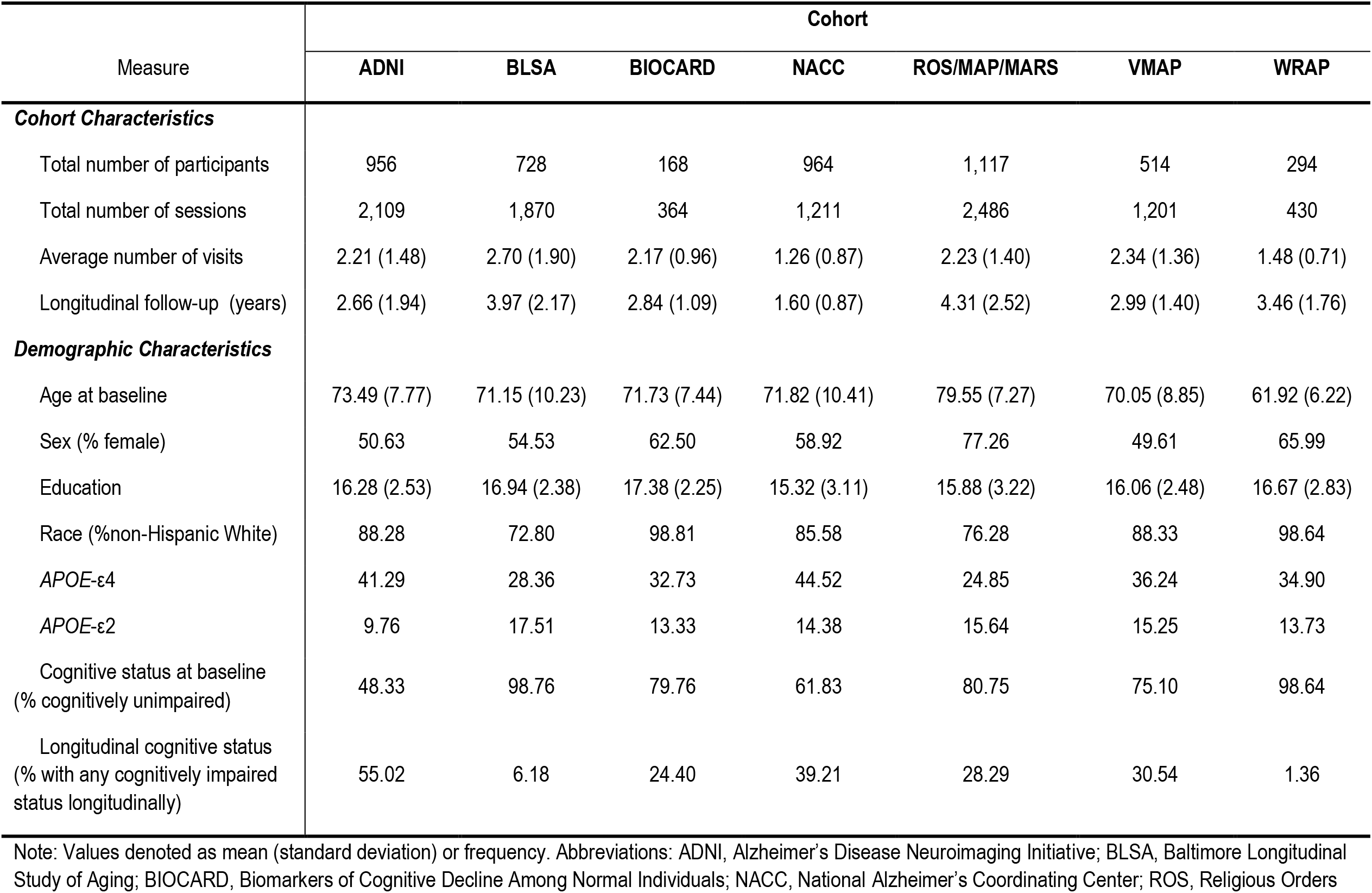
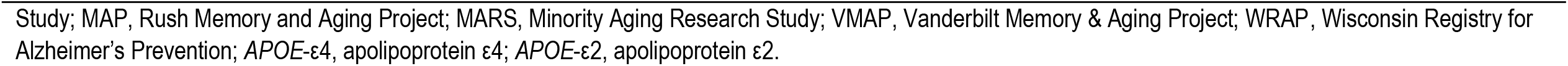
Demographic and health characteristics by cohort

| Measure | Cohort |  |  |  |  |  |  |
| --- | --- | --- | --- | --- | --- | --- | --- |
|  | ADNI | BLSA | BIOCARD | NACC | ROS/MAP/MARS | VMAP | WRAP |
| **Cohort Characteristics** |  |  |  |  |  |  |  |
| Total number of participants | 956 | 728 | 168 | 964 | 1,117 | 514 | 294 |
| Total number of sessions | 2,109 | 1,870 | 364 | 1,211 | 2,486 | 1,201 | 430 |
| Average number of visits | 2.21 (1.48) | 2.70 (1.90) | 2.17 (0.96) | 1.26 (0.87) | 2.23 (1.40) | 2.34 (1.36) | 1.48 (0.71) |
| Longitudinal follow-up (years) | 2.66 (1.94) | 3.97 (2.17) | 2.84 (1.09) | 1.60 (0.87) | 4.31 (2.52) | 2.99 (1.40) | 3.46 (1.76) |
| **Demographic Characteristics** |  |  |  |  |  |  |  |
| Age at baseline | 73.49 (7.77) | 71.15 (10.23) | 71.73 (7.44) | 71.82 (10.41) | 79.55 (7.27) | 70.05 (8.85) | 61.92 (6.22) |
| Sex (% female) | 50.63 | 54.53 | 62.50 | 58.92 | 77.26 | 49.61 | 65.99 |
| Education | 16.28 (2.53) | 16.94 (2.38) | 17.38 (2.25) | 15.32 (3.11) | 15.88 (3.22) | 16.06 (2.48) | 16.67 (2.83) |
| Race (%non-Hispanic White) | 88.28 | 72.80 | 98.81 | 85.58 | 76.28 | 88.33 | 98.64 |
| APOE-ε4 | 41.29 | 28.36 | 32.73 | 44.52 | 24.85 | 36.24 | 34.90 |
| APOE-ε2 | 9.76 | 17.51 | 13.33 | 14.38 | 15.64 | 15.25 | 13.73 |
| Cognitive status at baseline<br>(% cognitively unimpaired) | 48.33 | 98.76 | 79.76 | 61.83 | 80.75 | 75.10 | 98.64 |
| Longitudinal cognitive status<br>(% with any cognitively impaired<br>status longitudinally) | 55.02 | 6.18 | 24.40 | 39.21 | 28.29 | 30.54 | 1.36 |

Note: Values denoted as mean (standard deviation) or frequency. Abbreviations: ADNI, Alzheimer's Disease Neuroimaging Initiative; BLSA, Baltimore Longitudinal Study of Aging; BIOCARD, Biomarkers of Cognitive Decline Among Normal Individuals; NACC, National Alzheimer's Coordinating Center; ROS, Religious Orders
Study; MAP, Rush Memory and Aging Project; MARS, Minority Aging Research Study; VMAP, Vanderbilt Memory & Aging Project; WRAP, Wisconsin Registry for Alzheimer's Prevention; *APOE-ε4*, apolipoprotein ε4; *APOE-ε2*, apolipoprotein ε2.

### Diffusion MRI Acquisition and Preprocessing

All longitudinal dMRI data were preprocessed using the *PreQual* automated pipeline for denoising, slice-wise outlier imputation, data quality assurance, and to correct for motion, eddy current, and susceptibility-induced distortions.^45,46^ Following *PreQual* preprocessing, the cleaned data was input into DTIFIT. Simultaneously, the cleaned data was input into MATLAB code^27^ to calculate free-water (FW) and FW-corrected fractional anisotropy (FA_FWcorr_). Following map generation, we created a standard space representation by non-linearly registering (i.e., symmetric normalization and linear interpolation)^47^ the FA_CONV_ map to the FMRIB58_FA atlas. The resulting warp was applied to the FW and FA_FWcorr_ maps, which were used in all subsequent analyses. Imaging sessions of individuals with consistently large age-regressed outliers (more than 5 standard deviations from the mean) in white matter tract microstructural values were removed. All sample sizes reported in the manuscript have accounted for excluded participants.

### White Matter Tractography Templates

Consistent with prior work,^28,48-51^ we used freely accessible tractography templates (https://github.com/VUMC-VMAC/Tractography_Templates) to assess the white matter microstructure in our cohort (**Figure 1**). In total, we used 48 tract templates from seven different tract types, including association, limbic, projection, motor transcallosal, occipital transcallosal, parietal transcallosal, and prefrontal transcallosal areas.

**Figure 1.**
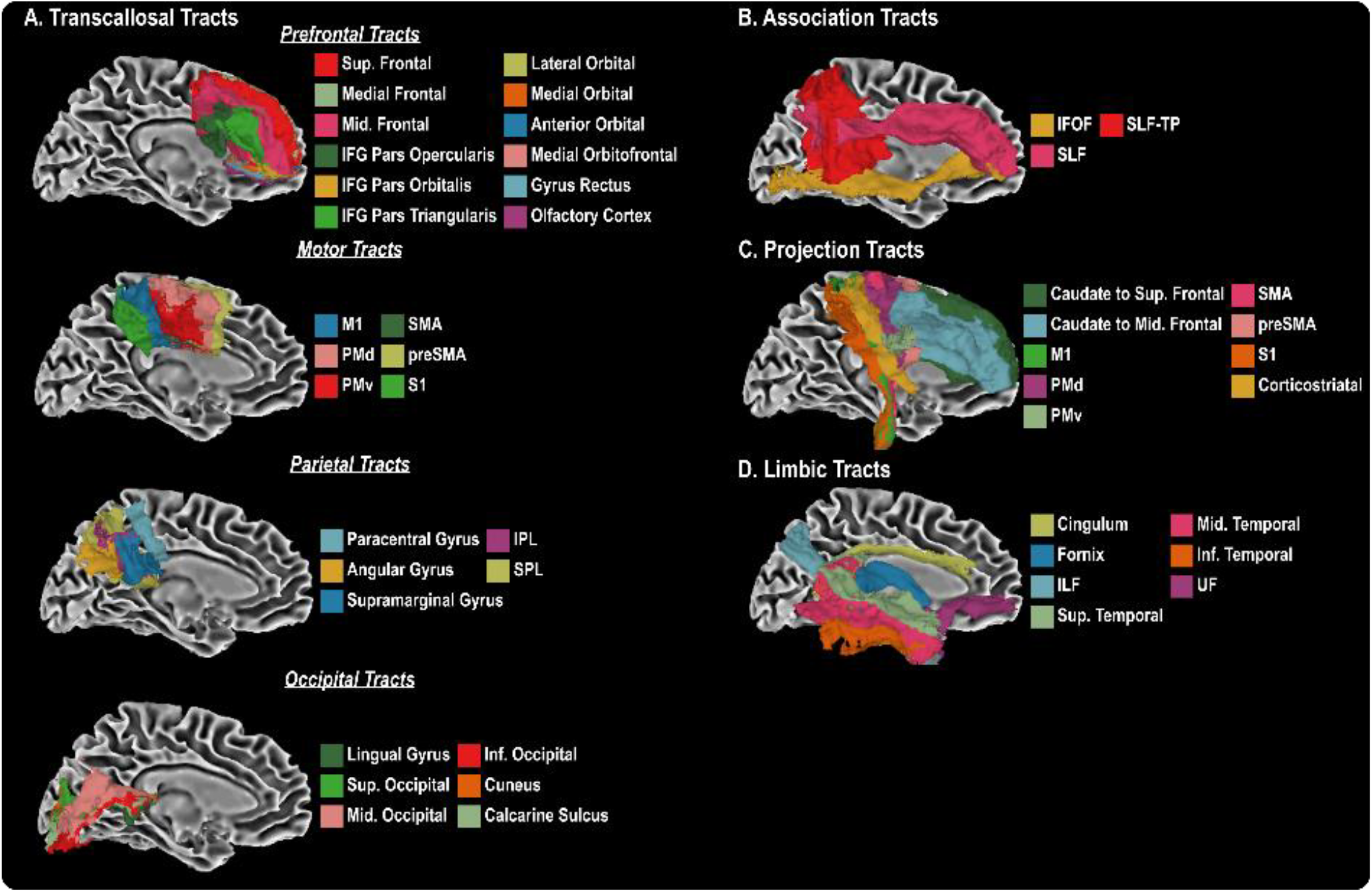
Forty-eight white matter tractography templates were used in the present study, and can be grouped into transcallosal (A), association (B), projection (C), and limbic tracts (D).

### Diffusion MRI Data Harmonization

A region-of-interest based approach was used to quantify mean FW and FA_FWcorr_ microstructural metrics across all tractography templates for each participant, resulting in a total of 96 unique values for each imaging session. These values were harmonized using the *Longitudinal ComBat* technique in R, controlling for all *site* × *scanner* × *protocol* combinations.^52^ A conservative harmonization approach was used to control for between-cohort effects, including mean-centered age, mean-centered age squared, sex, and diagnosis at baseline. We also included interactions of mean-centered age and converter status (i.e., cognitively unimpaired, cognitively impaired at some point) and mean-centered age squared and converter status. The harmonized values were then mean centered and used in all subsequent statistical analyses.

### Statistical Analyses

All statistical analyses were performed in R (version 4.1.0), and age was mean-centered prior to analysis. Covariates included age at baseline in addition to sex, race/ethnicity, and *APOE*-ε4 carrier status. Linear mixed-effects regression models were used to determine if sex, *APOE*-ε4 carrier status, or race were associated with FW and FA_FWcorr_ dMRI metrics over time. We controlled for the random aging effects for each participant (i.e., ∼1+age|participant). Models were repeated with *sex* × *APOE-*ε4, *sex* × *race, race* × *APOE-*ε4, and *sex* × *race x APOE-*ε4 interaction terms. Significance was set a priori as α=0.05 and corrected for multiple comparisons using the false discovery rate method.

## Results

### Demographic Differences

Females were more likely to be cognitively unimpaired at study entry (p=2.36×10^-23^) and less likely to develop cognitive impairment over time (p=1.94×10^-19^) compared to males. Non-Hispanic Black participants were more likely to be cognitively unimpaired at study entry (p=5.18×10^-5^) and longitudinally (p=5.03×10^-6^) compared to non-Hispanic White participants. *APOE-*ε4 carriers were more likely to be cognitively impaired at study entry (p=9.19×10^-31^) and more likely to develop cognitive impairment over time (p=7.31×10^-22^) compared to non-carriers. See **Supplemental Table 1** for differences across cohorts.

### Sex Associations with Longitudinal FA_FWcorr_ and FW

As illustrated in **Figure 2A**, females had lower FA_FWcorr_ over time relative to males across all 33 of the white matter tracts that showed a significant sex difference. **Table 2** summarizes the top five associations for sex, emphasizing the prominence of projection white matter tracts. Notably, the strongest association was observed in the ventral premotor projection tract, as depicted in the spaghetti plot within **Figure 2A** (p=3.87×10^-81^). **Figure 2B** illustrates the main effects of sex on FW, highlighting significant associations across many limbic, parietal TC, and occipital TC white matter tracts. In all 32 white matter tracts that showed a difference in FW by sex, females had a smaller increase in FW longitudinally relative to males. The strongest association was observed in the fornix, as depicted in **Figure 2B** (p=3.18×10^-23^). Statistics for all models can be found in **Supplemental Table 2**.

**Table 2.** Top Sex, Race, and *APOE*-ε4 Main Effects on Longitudinal White Matter Microstructure

| Tract | Tract Type | Outcome | β (SE) | z-value | p-value |
| --- | --- | --- | --- | --- | --- |
| **Main Effects of Sex** |  |  |  |  |  |
| PMv Projection | Projection | FA<sub>FWcorr</sub> | -9.82x10<sup>-3</sup> (5.08x10<sup>-4</sup>) | -19.31 | 3.87x10<sup>-81</sup> |
| S1 Projection | Projection | FA<sub>FWcorr</sub> | -9.13x10<sup>-3</sup> (6.86x10<sup>-4</sup>) | -13.31 | 9.46x10<sup>-39</sup> |
| SMA Projection | Projection | FA<sub>FWcorr</sub> | -7.36x10<sup>-3</sup> (5.60x10<sup>-4</sup>) | -13.15 | 5.75x10<sup>-38</sup> |
| M1 Projection | Projection | FA<sub>FWcorr</sub> | -7.69x10<sup>-3</sup> (6.04x10<sup>-4</sup>) | -12.72 | 1.01x10<sup>-35</sup> |
| PMd Projection | Projection | FA<sub>FWcorr</sub> | -6.90x10<sup>-3</sup> (5.43x10<sup>-4</sup>) | -12.71 | 1.01x10<sup>-35</sup> |
| **Main Effects of Race** |  |  |  |  |  |
| SLF-TP | Association | FA<sub>FWcorr</sub> | 6.59x10<sup>-3</sup> (7.29x10<sup>-4</sup>) | 9.04 | 1.22x10<sup>-17</sup> |
| preSMA Projection | Projection | FA<sub>FWcorr</sub> | 6.80x10<sup>-3</sup> (7.56x10<sup>-4</sup>) | 8.99 | 1.22x10<sup>-17</sup> |
| PMv Projection | Projection | FA<sub>FWcorr</sub> | 5.93x10<sup>-3</sup> (6.95x10<sup>-4</sup>) | 8.53 | 4.72x10<sup>-16</sup> |
| SMA Projection | Projection | FA<sub>FWcorr</sub> | 6.44x10<sup>-3</sup> (7.67x10<sup>-4</sup>) | 8.39 | 1.17x10<sup>-15</sup> |
| Supramarginal Gyrus TC | Parietal TC | FA<sub>FWcorr</sub> | 6.70x10<sup>-3</sup> (8.63x10<sup>-4</sup>) | 7.76 | 1.64x10<sup>-13</sup> |
| **Main Effects of APOE-ε4** |  |  |  |  |  |
| ILF | Limbic | FW | 6.70x10<sup>-3</sup> (9.99x10<sup>-4</sup>) | 6.71 | 1.03x10<sup>-9</sup> |
| Cingulum | Limbic | FW | 5.36x10<sup>-3</sup> (8.01x10<sup>-4</sup>) | 6.70 | 1.03x10<sup>-9</sup> |
| Inferior Temporal Gyrus | Limbic | FW | 5.58x10<sup>-3</sup> (9.34x10<sup>-4</sup>) | 5.97 | 7.47x10<sup>-8</sup> |
| Fornix | Limbic | FW | 6.99x10<sup>-3</sup> (1.22x10<sup>-3</sup>) | 5.73 | 2.36x10<sup>-7</sup> |
| Superior Occipital Gyrus TC | Occipital TC | FW | 6.32x10<sup>-3</sup> (1.13x10<sup>-3</sup>) | 5.61 | 3.99x10<sup>-7</sup> |

Abbreviations: FW, free-water; FA<sub>FWcorr</sub>, FW-corrected fractional anisotropy; ILF, inferior longitudinal fasciculus; PMd, dorsal premotor; PMv, ventral premotor; SLF-TP, superior longitudinal fasciculus-temporoparietal; SMA, supplementary motor area; TC, transcallosal. Listed p-values are corrected for multiple comparisons using the FDR approach.

**Figure 2.**
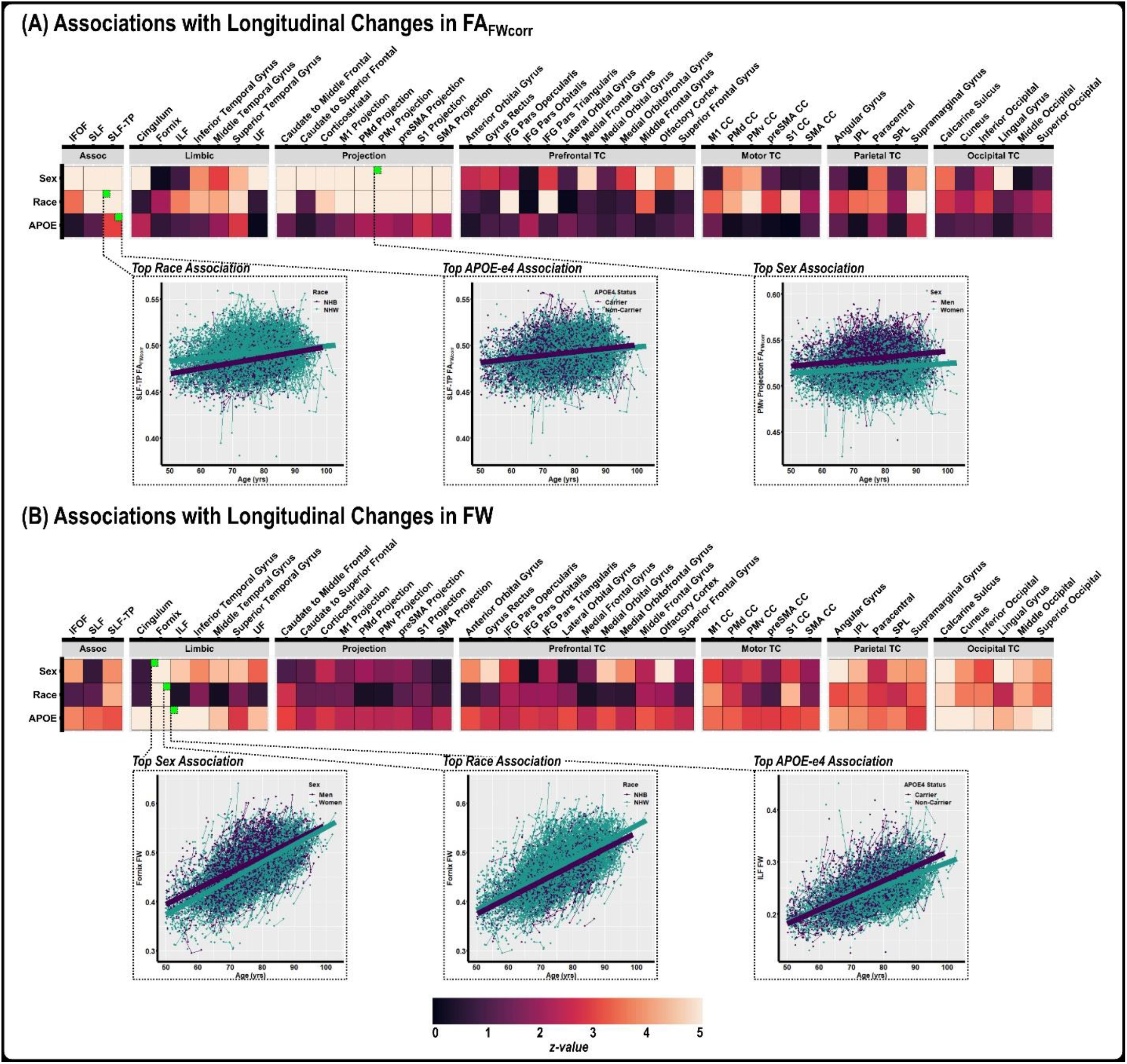
The main effects of sex, race, and *APOE*-ε4 on change in white matter microstructure. Linear mixed effects regression was conducted to determine the association of sex, race, and *APOE*-ε4 positivity on change in FA_FWcorr_ (**A**) and change in FW (**B**). For both measures, heatmaps are grouped by tract-type and illustrate the test statistic (i.e., z-value) for each independent regression analysis. The top race, sex, and *APOE*-ε4 associations for both measures are illustrated with spaghetti plots.

### Race Associations with Longitudinal FA_FWcorr_ and FW

Of the 32 tracts that were significantly different, non-Hispanic White adults had higher FA_FWcorr_ over time relative to non-Hispanic Black adults in all tracts except the fornix and inferior frontal-occipital fasciculus, as depicted in **Figure 2A**. The strongest association was observed in the superior longitudinal fasciculus-temporoparietal (SLF-TP) association tract, as depicted in the spaghetti plot within **Figure 2A** (p=1.22×10^-17^).**Table 2** highlights other strong associations, particularly within projection fibers. **Figure 2B** shows the main effects of race on FW. Of the 19 tracts that showed differences in FW by race, non-Hispanic White adults had lower FW values longitudinally relative to non-Hispanic Black adults in all tracts except the fornix. As seen in the spaghetti plot within Figure 2B, the strongest association was observed in the fornix (p=3.90×10^-12^) with higher fornix FW over time in non-Hispanic White adults relative to non-Hispanic Black adults. Statistics for all models can be found in **Supplemental Table 3**.

**Table 3.**
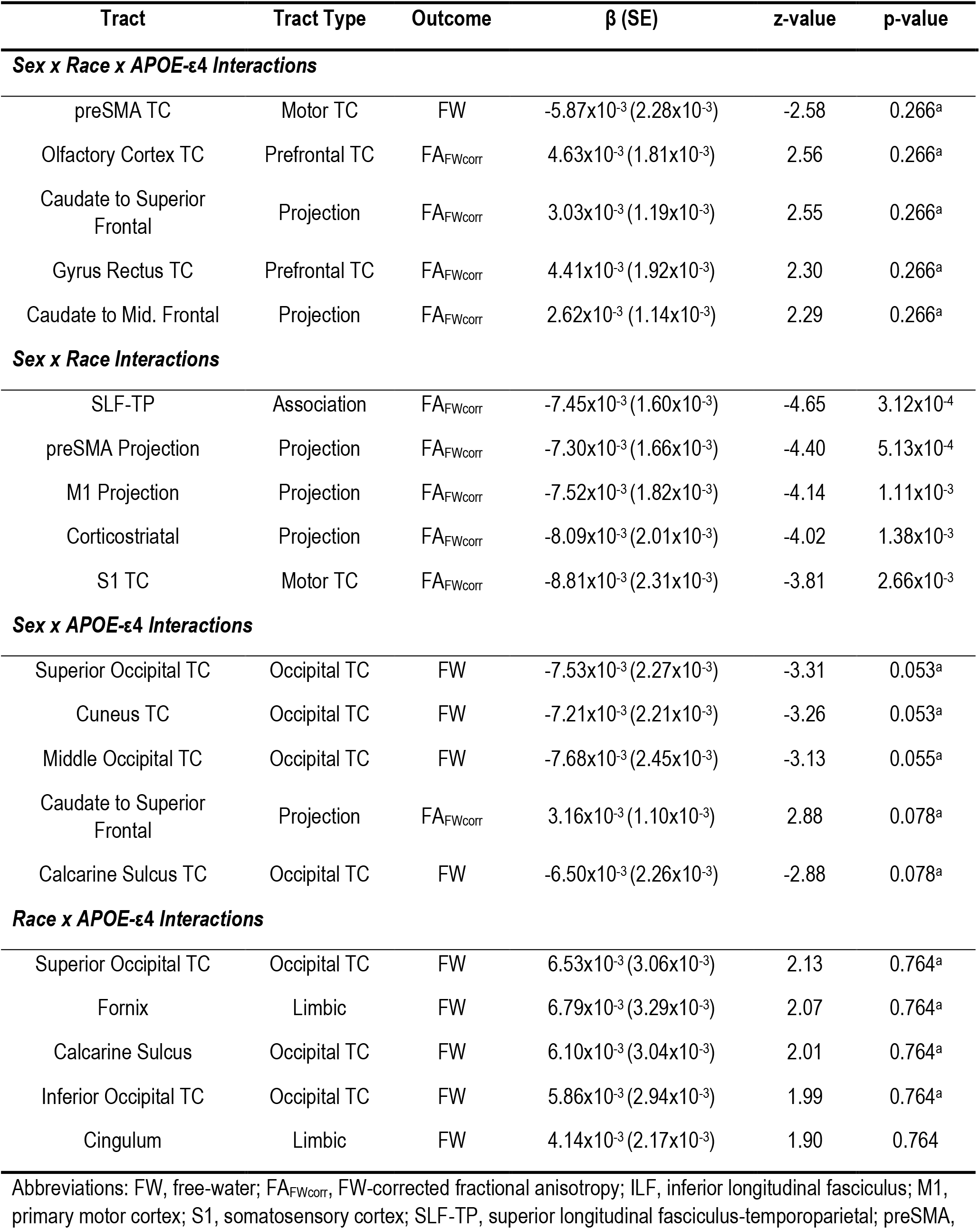
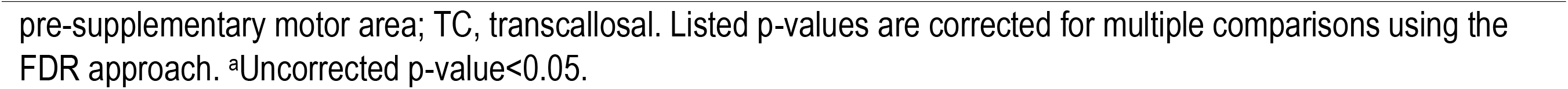
Top Sex, Race, and *APOE*-ε4 Interactions on Longitudinal White Matter Microstructure

| Tract | Tract Type | Outcome | β (SE) | z-value | p-value |
| --- | --- | --- | --- | --- | --- |
| **Sex x Race x APOE-ε4 Interactions** |  |  |  |  |  |
| preSMA TC | Motor TC | FW | -5.87x10<sup>-3</sup> (2.28x10<sup>-3</sup>) | -2.58 | 0.266<sup>a</sup> |
| Olfactory Cortex TC | Prefrontal TC | FA<sub>FWcorr</sub> | 4.63x10<sup>-3</sup> (1.81x10<sup>-3</sup>) | 2.56 | 0.266<sup>a</sup> |
| Caudate to Superior Frontal | Projection | FA<sub>FWcorr</sub> | 3.03x10<sup>-3</sup> (1.19x10<sup>-3</sup>) | 2.55 | 0.266<sup>a</sup> |
| Gyrus Rectus TC | Prefrontal TC | FA<sub>FWcorr</sub> | 4.41x10<sup>-3</sup> (1.92x10<sup>-3</sup>) | 2.30 | 0.266<sup>a</sup> |
| Caudate to Mid. Frontal | Projection | FA<sub>FWcorr</sub> | 2.62x10<sup>-3</sup> (1.14x10<sup>-3</sup>) | 2.29 | 0.266<sup>a</sup> |
| **Sex x Race Interactions** |  |  |  |  |  |
| SLF-TP | Association | FA<sub>FWcorr</sub> | -7.45x10<sup>-3</sup> (1.60x10<sup>-3</sup>) | -4.65 | 3.12x10<sup>-4</sup> |
| preSMA Projection | Projection | FA<sub>FWcorr</sub> | -7.30x10<sup>-3</sup> (1.66x10<sup>-3</sup>) | -4.40 | 5.13x10<sup>-4</sup> |
| M1 Projection | Projection | FA<sub>FWcorr</sub> | -7.52x10<sup>-3</sup> (1.82x10<sup>-3</sup>) | -4.14 | 1.11x10<sup>-3</sup> |
| Corticostriatal | Projection | FA<sub>FWcorr</sub> | -8.09x10<sup>-3</sup> (2.01x10<sup>-3</sup>) | -4.02 | 1.38x10<sup>-3</sup> |
| S1 TC | Motor TC | FA<sub>FWcorr</sub> | -8.81x10<sup>-3</sup> (2.31x10<sup>-3</sup>) | -3.81 | 2.66x10<sup>-3</sup> |
| **Sex x APOE-ε4 Interactions** |  |  |  |  |  |
| Superior Occipital TC | Occipital TC | FW | -7.53x10<sup>-3</sup> (2.27x10<sup>-3</sup>) | -3.31 | 0.053<sup>a</sup> |
| Cuneus TC | Occipital TC | FW | -7.21x10<sup>-3</sup> (2.21x10<sup>-3</sup>) | -3.26 | 0.053<sup>a</sup> |
| Middle Occipital TC | Occipital TC | FW | -7.68x10<sup>-3</sup> (2.45x10<sup>-3</sup>) | -3.13 | 0.055<sup>a</sup> |
| Caudate to Superior Frontal | Projection | FA<sub>FWcorr</sub> | 3.16x10<sup>-3</sup> (1.10x10<sup>-3</sup>) | 2.88 | 0.078<sup>a</sup> |
| Calcarine Sulcus TC | Occipital TC | FW | -6.50x10<sup>-3</sup> (2.26x10<sup>-3</sup>) | -2.88 | 0.078<sup>a</sup> |
| **Race x APOE-ε4 Interactions** |  |  |  |  |  |
| Superior Occipital TC | Occipital TC | FW | 6.53x10<sup>-3</sup> (3.06x10<sup>-3</sup>) | 2.13 | 0.764<sup>a</sup> |
| Fornix | Limbic | FW | 6.79x10<sup>-3</sup> (3.29x10<sup>-3</sup>) | 2.07 | 0.764<sup>a</sup> |
| Calcarine Sulcus | Occipital TC | FW | 6.10x10<sup>-3</sup> (3.04x10<sup>-3</sup>) | 2.01 | 0.764<sup>a</sup> |
| Inferior Occipital TC | Occipital TC | FW | 5.86x10<sup>-3</sup> (2.94x10<sup>-3</sup>) | 1.99 | 0.764<sup>a</sup> |
| Cingulum | Limbic | FW | 4.14x10<sup>-3</sup> (2.17x10<sup>-3</sup>) | 1.90 | 0.764 |

Abbreviations: FW, free-water; FA<sub>FWcorr</sub>, FW-corrected fractional anisotropy; ILF, inferior longitudinal fasciculus; M1, primary motor cortex; S1, somatosensory cortex; SLF-TP, superior longitudinal fasciculus-temporoparietal; preSMA,
pre-supplementary motor area; TC, transcallosal. Listed p-values are corrected for multiple comparisons using the FDR approach. <sup>a</sup>Uncorrected p-value<0.05.

### APOE-ε4 Associations with Longitudinal FA_FWcorr_ and FW

**Figure 2A** illustrates that the effects of *APOE-*ε4 on FA_FWcorr_ were primarily limited to select limbic and motor TC fibers. Eight tracts showed differences by *APOE-*ε4 status, with *APOE-*ε4 carriers having higher FA_FWcorr_ longitudinally in all tracts except the inferior frontal gyrus pars opercularis prefrontal transcallosal tract. The strongest association was observed in the SLF-TP association tract, as depicted in the spaghetti plot within **Figure 2A** (p=4.82×10^-3^). FW was greater over time in *APOE-*ε4 carriers in all tracts examined except the primary somatosensory cortex (**Figure 2B**) As shown in **Table 2**, the strongest associations were observed for FW in limbic and occipital TC fibers. Statistics for all models can be found in **Supplemental Table 4**.

### Sex, Race, and *APOE-*ε4 Interactions with Longitudinal FA_FWcorr_ and FW

As summarized in **Table 3**, there were no statistically significant sex × *APOE-*ε4, race × *APOE-* ε4, or sex × race × *APOE-*ε4 interactions on FA_FWcorr_ or FW when corrected for multiple comparisons. Notably, race interacted with sex on FA_FWcorr_, primarily in projection fibers, though the strongest interaction was observed in the SLF-TP (p=3.12×10^-4^). Statistics for all models can be found in **Supplemental Tables 5-8**.

## Discussion

This study leveraged nine well-defined cohorts of older adults to characterize the effects of sex, race, and *APOE-*ε4 carrier status on the white matter integrity of 48 tracts longitudinally using FW and FA_FWcorr_ metrics. We found that sex and race were most strongly associated with change in FA_FWcorr_, whereby females and non-Hispanic Black adults had greater microstructural damage over time than males and non-Hispanic white adults. Sex differences were greatest in association and projection tracts, while race was associated primarily with projection tracts. *APOE*-ε4 status was strongly associated with FW in limbic and occipital TC tracts such that *APOE*-ε4 carriers had greater extracellular water than non-carriers over time. *APOE-*ε4 status did not modify the relationships between sex or race on FW or FA_FWcorr_, but race did modify the association between sex and FA_FWcorr_ in some projection fibers. This work adds to the field by exploring differences in white matter by sex, race, and *APOE*-ε4 carrier status on a larger scale than previously done.

In the present study we found that females had greater microstructural tissue damage than males longitudinally, and that these differences were most pronounced in association and projection tracts. This work advances the field by showing clear sex differences in white matter integrity. Projection tracts connect the cortex to subcortical structures and help integrate sensory, motor, and cognitive functions.Our findings support previous cross-sectional work that found that females had lower FA values in sensory and motor projections^10^ and a greater decline in FA over time compared to males.^11^ Previous work has consistently found that association tracts are affected in the aging process^10,28^ and these tracts show a greater decline in FA and FA_FWcorr_ in individuals with cognitive impairment relative to cognitively unimpaired adults.^28,53^ While some studies have shown that projection fibers are also associated with abnormal cognitive aging, these associations are less robust.^10,28,49^ Given that females are disproportionately affected by AD, the observed sex differences in FA_FWcorr_ in our study suggest that white matter integrity may contribute to this disparity. However, we also found that in tracts with sex differences in FW, females tended to have lower values than males longitudinally. Although FW has been proposed to have many possible neurobiological correlates, including atrophy, edema, and neuroinflammation, the etiology of extracellular water is not agreed upon.^54^ We found that females in this study were less likely to be cognitively impaired at baseline and longitudinally than males, which could explain why they had lower FW values than males. Future studies to determine whether observed changes in FW are compensatory and beneficial are necessary.

We showed that longitudinally non-Hispanic White adults tended to have higher FA_FWcorr_ than non-Hispanic Black adults. These differences were most pronounced in the SLF, some transcallosal tracks, and projection tracts. Small cross-sectional studies of Black women^55,56^ have shown that racial discrimination is associated with worse white matter integrity in the corpus callosum, cingulum, and SLF, tracts known to be affected in AD. Studies comparing racial and ethnic groups have suggested that other social factors, including socioeconomic status^57^ and acculturation,^58^ also affect white matter integrity. We believe the observed racial differences in this study relate to sociocultural factors as opposed to biological differences between racial groups. Additional work to explore possible mediators of the relationship between social determinants of health and white matter integrity, such as vascular risk factors, are warranted. We found fewer differences in FW. While some studies suggest that associations with FA are driven by FW,^54^ we found independent effects, suggesting FW and FW-corrected measures provide complementary information about biological processes related to white matter integrity. In tracts that did have differences in FW, non-Hispanic White adults tended to have lower FW than non-Hispanic Black adults. If differences in microstructure were due to AD pathology alone, we might expect non-Hispanic Black adults in this study to have better microstructure as they were less likely to be cognitively impaired at baseline and longitudinally compared to non-Hispanic White adults. That we did not further suggests that there are other factors, including social factors, that affect white matter integrity. Interestingly, in the fornix FW was elevated and FA_FWcorr_ was lower in non-Hispanic White participants relative to non-Hispanic Black participants. Because the fornix is adjacent to the ventricles, it is particularly prone to partial volume effects,^27,59^ and additional studies assessing drivers of microstructure of the fornix are warranted. Race modified the association between sex and FA_FWcorr_, primarily in projection tracts. One possible explanation is that due to intersectionality (e.g., intersecting identities), non-Hispanic Black women may experience discrimination on the basis of both race and gender,^60^ driving increased white matter deterioration, though further work to understand this effect modification is necessary.

*APOE*-ε4 carrier status was associated with elevated FW, primarily in limbic and occipital transcallosal tracts. There were fewer differences in FA_FWcorr_. Despite the role of *APOE* in lipid homeostasis, studies do not consistently find an association between *APOE*-ε4 and worse white matter integrity.^61-63^ In previous work, we have shown that the FW measure, particularly in limbic tracts, is sensitive to abnormal aging.^28,49^ This study extends this work by showing that *APOE*-ε4 carriers, who are known to be at greater risk for AD, show these same changes in the FW metric in limbic tracts. This suggests that processes that increase interstitial spaces, such as atrophy, inflammation, or edema, are present to a greater extent in *APOE*-ε4 carriers in tracts known to be affected in AD. Given that neither sex nor race moderated the association between *APOE*-ε4 and FW, observed differences in *APOE*-ε4 and AD risk by sex and race may not be driven by the same pathological processes that drives FW changes.

This study has several strengths. It used a large harmonized multi-site diffusion MRI cohort, looking at 48 tractography templates that included association, limbic, projection, and transcallosal tracts, allowing us to explore sex, racial, and *APOE*-ε4 carrier differences in white matter integrity. Additionally, we were able to account for partial volume effects by using FW and FW-corrected diffusion metrics. This study also had limitations. While non-Hispanic Black participants were included, they were primarily drawn from two cohorts and were less likely to be cognitively impaired at baseline and longitudinally than non-Hispanic White participants, which is the opposite of what epidemiological studies show.

Differences in cohort demographics may contribute to observed differences in white matter integrity and could temper the conclusions that can be drawn. Additionally, although this study found notable differences in white matter integrity, we are not able to determine the biological and social pathways that lead to these differences.

In this study we demonstrate marked differences longitudinally in white matter integrity by sex, race and *APOE*-ε4 carrier status on a larger scale than previously done. Females, non-Hispanic Black adults, and *APOE*-ε4 carriers had measures suggestive of worse white matter integrity compared to males, non-Hispanic White adults, and *APOE*-ε4 non-carriers. The markers of white matter integrity and the tracts affected were not uniform, suggesting there may be different etiologies for the observed differences. Future studies that incorporate larger samples sizes of diverse participants, additional biomarkers, and information about social determinants of health are needed to clarify the reasons for the observed differences.

## Notes

**Conflicts of Interest and Disclosure Statement:** SCJ has served on advisory boards for Enigma Biomedical and ALZPath in the past two years. AJS receives support from multiple NIH grants (P30 AG010133, P30 AG072976, R01 AG019771, R01 AG057739, U19 AG024904, R01 LM013463, R01 AG068193, T32 AG071444, U01 AG068057, U01 AG072177, U19 AG074879, and U24 AG074855). He has also received support from Avid Radiopharmaceuticals, a subsidiary of Eli Lilly (in kind contribution of PET tracer precursor) and participated in Scientific Advisory Boards (Bayer Oncology, Eisai, Novo Nordisk, and Siemens Medical Solutions USA, Inc) and an Observational Study Monitoring Board (MESA, NIH NHLBI), as well as External Advisory Committees for multiple NIA grants. He also serves as Editor-in-Chief of Brain Imaging and Behavior, a Springer-Nature Journal.

**Funding Acknowledgements:** This study was supported by several funding sources, including K01-EB032898 (KGS), R01-EB017230 (BAL) K01-AG073584 (DBA), U24-AG074855 (TJH), 75N95D22P00141 (TJH), R01-AG059716 (TJH), UL1-TR000445 and UL1-TR002243 (Vanderbilt Clinical Translational Science Award), S10-OD023680 (Vanderbilt’s High-Performance Computer Cluster for Biomedical Research). The research was support in part by the Intramural Research Program of the National Institutes of Health, National Institute on Aging. Study data were obtained from the Vanderbilt Memory and Aging Project (VMAP). VMAP data were collected by Vanderbilt Memory and Alzheimer’s Center Investigators at Vanderbilt University Medical Center. This work was supported by NIA grants R01-AG034962 (PI: Jefferson), R01-AG056534 (PI: Jefferson), R01-AG062826 (PI: Gifford), U19-AG03655 (PI:Albert) and Alzheimer’s Association IIRG-08-88733 (PI: Jefferson). The data contributed from the Wisconsin Registry for Alzheimer’s Prevention was supported by NIA AG021155, AG0271761, AG037639, and AG054047. The BLSA is supported by the Intramural Research Program of the National Institutes of Health, National Institute on AgingThis research was supported in part by the Intramural Research Program of the National Institutes of Health, National Institute on Aging. Data collection and sharing for this project was funded (in part) by the Alzheimer’s Disease Neuroimaging Initiative (ADNI) (National Institutes of Health Grant U01 AG024904) and DOD ADNI (Department of Defense award number W81XWH-12-2-0012). ADNI is funded by the National Institute on Aging, the National Institute of Biomedical Imaging and Bioengineering, and through generous contributions from the following: AbbVie, Alzheimer’s Association; Alzheimer’s Drug Discovery Foundation; Araclon Biotech; BioClinica, Inc.; Biogen; Bristol-Myers Squibb Company; CereSpir, Inc.; Cogstate; Eisai Inc.; Elan Pharmaceuticals, Inc.; Eli Lilly and Company; EuroImmun; F. Hoffmann-La Roche Ltd and its affiliated company Genentech, Inc.; Fujirebio; GE Healthcare; IXICO Ltd.; Janssen Alzheimer Immunotherapy Research & Development, LLC.; Johnson & Johnson Pharmaceutical Research & Development LLC.; Lumosity; Lundbeck; Merck & Co., Inc.; Meso Scale Diagnostics, LLC.; NeuroRx Research; Neurotrack Technologies; Novartis Pharmaceuticals Corporation; Pfizer Inc.; Piramal Imaging; Servier; Takeda Pharmaceutical Company; and Transition Therapeutics. The Canadian Institutes of Health Research is providing funds to support ADNI clinical sites in Canada. Private sector contributions are facilitated by the Foundation for the National Institutes of Health (www.fnih.org). The grantee organization is the Northern California Institute for Research and Education, and the study is coordinated by the Alzheimer’s Therapeutic Research Institute at the University of Southern California. ADNI data are disseminated by the Laboratory for Neuro Imaging at the University of Southern California. Additional acknowledgments include the National Institute on Aging Genetics of Alzheimer’s Disease Data Storage Site (NIAGADS, U24AG041689) at the University of Pennsylvania, funded by NIA. Data contributed from MAP/ROS/MARS was supported by NIA R01AG017917, P30AG10161, P30AG072975, R01AG022018, R01AG056405, UH2NS100599, UH3NS100599, R01AG064233, R01AG15819 and R01AG067482, and the Illinois Department of Public Health (Alzheimer’s Disease Research Fund). Data can be accessed at www.radc.rush.edu. The NACC database is funded by NIA/NIH Grant U24 AG072122. NACC data are contributed by the NIA-funded ADCs : P50 AG005131 (PI James Brewer, MD, PhD), P50 AG005133 (PI Oscar Lopez, MD), P50 AG005134 (PI Bradley Hyman, MD, PhD), P50 AG005136 (PI Thomas Grabowski, MD), P50 AG005138 (PI Mary Sano, PhD), P50 AG005142 (PI Helena Chui, MD), P50 AG005146 (PI Marilyn Albert, PhD), P50 AG005681 (PI John Morris, MD), P30 AG008017 (PI Jeffrey Kaye, MD), P30 AG008051 (PI Thomas Wisniewski, MD), P50 AG008702 (PI Scott Small, MD), P30 AG010124 (PI John Trojanowski, MD, PhD), P30 AG010129 (PI Charles DeCarli, MD), P30 AG010133 (PI Andrew Saykin, PsyD), P30 AG010161 (PI David Bennett, MD), P30 AG012300 (PI Roger Rosenberg, MD), P30 AG013846 (PI Neil Kowall, MD), P30 AG013854 (PI Robert Vassar, PhD), P50 AG016573 (PI Frank LaFerla, PhD), P50 AG016574 (PI Ronald Petersen, MD, PhD), P30 AG019610 (PI Eric Reiman, MD), P50 AG023501 (PI Bruce Miller, MD), P50 AG025688 (PI Allan Levey, MD, PhD), P30 (PI Linda Van Eldik, PhD), P50 AG033514 (PI Sanjay Asthana, MD, FRCP), P30 AG035982 (PI Russell Swerdlow, MD), P50 AG047266 (PI Todd Golde, MD, PhD), P50 AG047270 (PI Stephen Strittmatter, MD, PhD), P50 AG047366 (PI Victor Henderson, MD, MS), P30 AG049638 (PI Suzanne Craft, PhD), P30 AG053760 (PI Henry Paulson, MD, PhD), P30 AG066546 (PI Sudha Seshadri, MD), P20 AG068024 (PI Erik Roberson, MD, PhD), P20 AG068053 (PI Marwan Sabbagh, MD), P20 AG068077 (PI Gary Rosenberg, MD), P20 AG068082 (PI Angela Jefferson, PhD), P30 AG072958 (PI Heather Whitson, MD), P30 AG072959 (PI James Leverenz, MD). NACC data can be accessed at naccdata.org.

### Competing Interest Statement

SCJ has served on advisory boards for Enigma Biomedical and ALZPath in the past two years. AJS receives support from multiple NIH grants (P30 AG010133, P30 AG072976, R01 AG019771, R01 AG057739, U19 AG024904, R01 LM013463, R01 AG068193, T32 AG071444, U01 AG068057, U01 AG072177, U19 AG074879, and U24 AG074855). He has also received support from Avid Radiopharmaceuticals, a subsidiary of Eli Lilly (in kind contribution of PET tracer precursor) and participated in Scientific Advisory Boards (Bayer Oncology, Eisai, Novo Nordisk, and Siemens Medical Solutions USA, Inc) and an Observational Study Monitoring Board (MESA, NIH NHLBI), as well as External Advisory Committees for multiple NIA grants. He also serves as Editor-in-Chief of Brain Imaging and Behavior, a Springer-Nature Journal.

